# XenoType: Decoding cellular identity in breast cancer histology through molecular supervision

**DOI:** 10.64898/2026.11.24.768971

**Authors:** Duarte F. Forjaz, Diogo Fróis Vieira, Vasco Queiroga, Marvin Willig, Saurabh Joshi, Ashley L. Kiemen, André Forjaz, Denis Wirtz, Pei-Hsun Wu

**Author notes:** These authors contributed equally (Duarte F. Forjaz, Diogo Fróis Vieira, Vasco Queiroga, Marvin Willig). Send correspondence to: Denis Wirtz, Ph.D. 3400 N Charles St, Croft Hall 130, Baltimore MD, 21218, Pei-Hsun Wu, Ph.D. 3400 N Charles St, Croft Hall 130, Baltimore MD, 21218.

## Abstract

Characterizing the cellular composition of the tumor microenvironment from histological images is fundamental to understanding cancer biology and informing treatment decisions. Deep learning has enabled automated nuclear detection and classification from hematoxylin and eosin (H&E) images, yet cell classifiers remain largely confined to broad morphological categories owing to a reliance on manual annotations. Recent efforts have used spatial molecular measurements to supervise histological classification, but these approaches often require separate tissue sections or produce altered H&E staining, and the resulting cell-type annotations depend on subjective choices such as marker selection and clustering parameters. Here we develop XenoType, a framework that classifies cells into seven molecularly informed categories directly from routine histology, trained using paired same-section Xenium spatial transcriptomics. Cell-type annotations are reconciled from four single-cell RNA-sequencing references, reducing dependence on marker choices. Across eight held-out breast tissue sections, XenoType outperformed commonly used CellViT++ retrained on the same molecular annotations under both four-class (macro-F1: 0.703 vs. 0.569) and seven-class (macro-F1: 0.571 vs. 0.403) taxonomies, with the largest gains for myoepithelial (F1: 0.175 to 0.572) and endothelial cells (F1: 0.395 to 0.682). Molecular measurements are required only during training; at inference, XenoType uses the H&E image alone. Model-reference agreement increased consistently with annotation confidence, and XenoType generalized to an independently profiled breast tissue section from a different spatial transcriptomics platform, supporting scalable cell-type mapping from routine histology without per-specimen molecular profiling.

## Introduction

Breast cancer comprises heterogeneous populations of malignant and nonmalignant cells whose composition and spatial organization influence disease progression, immune evasion and clinical outcome^1–4^. Hematoxylin and eosin (H&E) staining provides a detailed view of tissue architecture and remains central to histopathological assessment^5,6^. However, morphologically similar cells may exhibit distinct molecular identities, limiting the cellular resolution achievable through conventional histological examination^7^. Resolving these populations and their spatial relationships directly from H&E images would enable more detailed characterization of tumor heterogeneity across routinely collected tissue specimens and the large collections already available in clinical archives^8–11^.

Deep learning has enabled automated nuclear detection and classification in histological images, with recent foundation models providing transferable representations of tissue morphology^12–14^. Nevertheless, cell classifiers remain constrained by training data: existing annotated datasets rely predominantly on manual labels, which are labor-intensive, subject to observer variability, and generally restricted to broad morphological categories, including the human breast^15,16^. Spatial transcriptomics offers an alternative source of supervision by measuring gene expression while preserving the spatial organization at cellular or tissue resolution^2,17–21^. Recent studies have begun to address these limitations by leveraging molecular measurements as supervision for histological analysis^18,22^. Li *et al.* used co-registered multiplex immunofluorescence to generate protein-based cell annotations for training H&E classifiers^22^, while STHELAR established a multi-tissue resource linking spatial transcriptomic measurements with histological images^18^. Other frameworks have extended this direction toward predicting spatially resolved gene expression and cell identities directly from H&E^23–25^.

Although these approaches demonstrate the potential of molecularly supervised classification, important gaps remain. Many spatial molecular assays, such as multiplex immunofluorescence, either require a separate tissue section or alter the H&E stain, since staining is typically performed after molecular profiling rounds^26,27^. In addition, cell-type assignments in spatial transcriptomics datasets are typically assigned by clustering expression profiles and manually inspecting marker genes for each cluster - a process that is subjective, difficult to reproduce and sensitive to the choice of markers and resolution parameters^28,29^. Automated reference-based methods can reduce this subjectivity by transferring labels from annotated single-cell datasets. However, the resulting assignments depend on references that differ in cellular composition and annotation granularity, and systematic strategies to reconcile predictions across references and to characterize annotation concordance remain limited^30,31^. At the model level, leading frameworks for cell analysis derive classification features from their segmentation backbone, coupling the two tasks and restricting the independent selection of representations best suited for distinguishing finer cellular populations^12–14,32^. Together, these constraints have kept histological cell classification largely confined to broad morphological categories and motivate a framework that combines molecularly informed annotations with flexible histological feature extraction^15,16,22^.

Here, we introduce XenoType, a framework for training cell classifiers on H&E images using annotations derived from paired Xenium spatial transcriptomics, where H&E staining is performed on the same tissue section as molecular profiling. We reconcile cell-type predictions from four single-cell RNA-sequencing references into seven categories and transfer the resulting labels to nuclei detected in H&E. XenoType separates nuclear segmentation from feature extraction, allowing a pathology foundation model to supply cell representations while retaining the same segmentation model. XenoType achieved higher agreement with molecular reference annotations than CellViT++ retrained on the same labels across eight held-out Xenium breast tissue sections and an additional section profiled using ATERA. At inference, XenoType requires only the H&E image, supporting scalable cell-type mapping across tissue collections without molecular profiling of each specimen.

## Results

### Spatial transcriptomics enables molecular supervision of histological cell classification

We developed XenoType to classify individual cells in images of standard hematoxylin and eosin (H&E)-stained tissue sections using molecularly informed training annotations (**Fig. 1**). The study used paired Xenium spatial transcriptomic and histological data from the same tissue section of 12 publicly available human breast tissue specimens, linking gene expression profiles to tissue morphology at single-cell resolution. Using four independent single-cell RNA-sequencing references (Yan, Wu, Dong, and Chen)^2,33–35^, we assigned Xenium cells to seven categories: tumor, epithelial, myoepithelial, endothelial, myeloid, lymphocyte, and stromal cells. We transferred these labels to nuclei detected in the corresponding H&E images, establishing a training resource that associates histological morphology with molecularly inferred cell identities. Four tissue sections were selected for model development to capture diverse histological appearances and cellular populations, while the remaining eight were reserved for independent evaluation under four-class (PanNuke-aligned) and seven-class classification schemes^15^.

**Figure 1.**
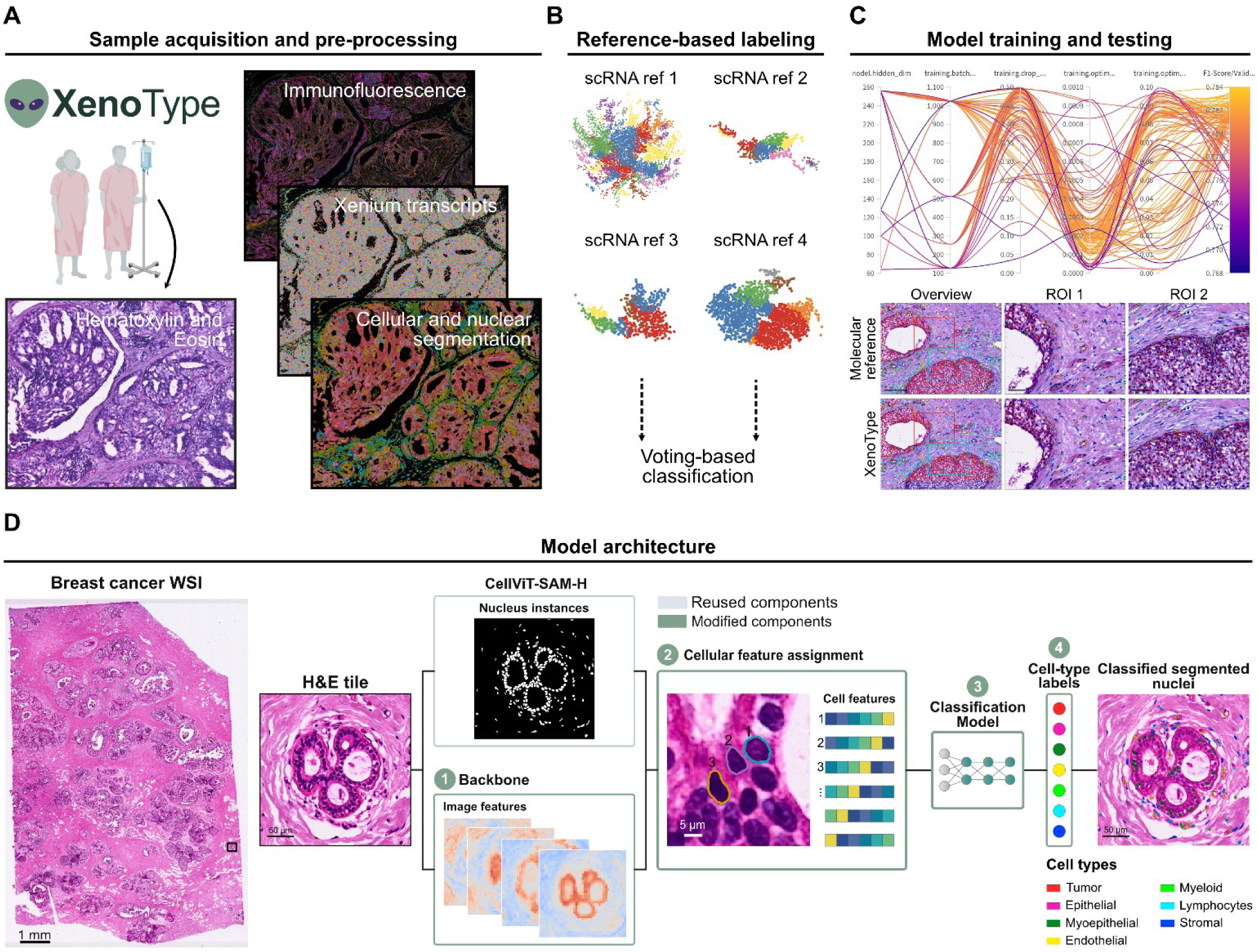
XenoType infers cell classification from standard H&E images. **(A)** Paired hematoxylin and eosin (H&E) imaging, immunofluorescence imaging, and Xenium spatial transcriptomic profiling are performed on the same human breast tissue section. **(B)** Xenium cells are independently annotated using four single-cell RNA-sequencing (scRNA-seq) reference datasets. Reference predictions are harmonized into seven cellular categories and reconciled through voting and class-specific annotation rules. **(C)** A parallel coordinates plot (top) summarizes the configuration search across feature extractors, classifier heads, and training hyperparameters, with lines colored by validation F1 score. Representative tissue regions (bottom) compare molecular reference annotations and XenoType predictions at overview and enlarged scales. **(D)** XenoType separates nuclear segmentation from foundation-model feature extraction for histological cell classification. H&E tiles are processed through parallel nuclear segmentation (CellViT-SAM-H) and feature-extraction (H-Optimus-1) branches. Cellular feature assignment associates these representations with individual segmented nuclei to generate cell-level embeddings. A classification head predicts cellular identities from the resulting features, which are combined with nuclear instance masks to generate spatially resolved classification maps. Gray and turquoise indicate reused and modified components, respectively. Output colors identify the seven molecularly informed cell categories.

### Multi-reference annotation defines seven molecularly informed cell categories

Spatial transcript density and cell-level gene expression profiles summarize the molecular data used for annotation (**Fig. 2A**). The four reference datasets^2,33–35^ differed in cellular composition and class representation, as illustrated by their respective embeddings (**Fig. 2B**). We used SingleR^36^ to annotate each Xenium cell independently against each reference, then mapped the reference-specific predictions to the seven shared categories.

**Figure 2.**
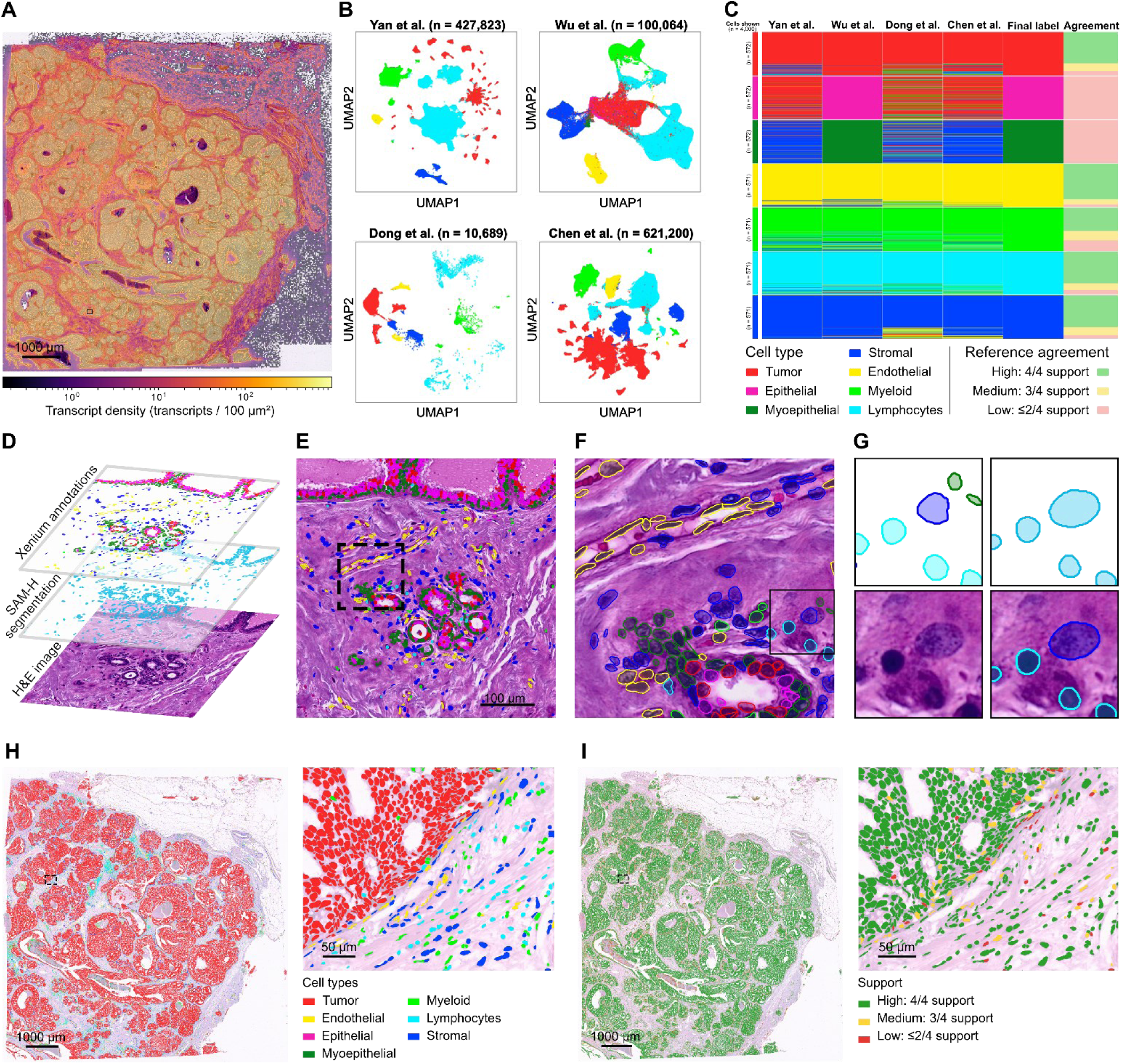
Multi-reference transcriptomic annotation defines seven cell categories and reveals differences in reference agreement. **(A)** Spatial transcript density overlaid on a representative breast tissue H&E section. Color scale indicates transcripts per 100 μm2. **(B)** Uniform manifold approximation and projection (UMAP) embeddings of the four single-cell RNA-sequencing (scRNA-seq) reference datasets (Yan, Wu, Dong, and Chen), colored by harmonized cell category. **(C)** Reference-specific cell-type predictions, final reconciled labels, and supporting-reference counts for 4,000 cells sampled across the seven categories. Epithelial and myoepithelial assignments incorporate a class-specific rule based on the Wu referencewhere applicable. **(D)** Spatial correspondence between Xenium annotations, CellViT-SAM-H nuclear segmentation, and the underlying H&E image in a representative tissue region. **(E)** Region shown in D with Xenium boundaries and cell classifications overlaid on H&E. **(F)** Enlarged region (dashed box in E) illustrating individual nuclear morphology at cellular scale. **(G)** Further enlargement of box in F showing incorrectly segmented Xenium boundaries (top left), correct SAM-H segmentation (top right), corresponding H&E region (bottom left), and SAM -H segmentation with classification transferred from Xenium overlaid on H&E (bottom right). **(H)** Spatial distribution of the seven annotated cell populations in a representative breast tissue section (left) and an enlarged region (right). **(I)** Corresponding spatial maps of reference agreement, categorized as high (4/4 supporting references), medium (3/4), or low (≤2/4). Reference agreement describes concordance among computational annotations rather than independently validated cell-type accuracy.

We reconciled the four SingleR predictions by voting and resolved ties using the mean SingleR score and a predefined reference order. Given only one reference (Wu^2^) had normal epithelial and myoepithelial cells, this reference’s assignment of these classes superseded other references’ votes. For each cell, we recorded how many references supported the final assignment. The 4,000-cell sample illustrates these patterns across the seven categories (**Fig. 2C**). Because reference datasets differed in the categories they contained, limited support for epithelial and myoepithelial labels partly reflects reference coverage. The Wu dataset was the only reference with labels that mapped to the epithelial and myoepithelial categories. Assignments to either category could therefore receive support from at most one of the four references. The spatial maps show the distribution of assigned categories and their supporting reference counts (**Fig. 2H,I**).

Across the 12 sections, 3,316,221 Xenium cells met the minimum thresholds of 10 transcripts and five detected genes per cell and were assigned molecular labels. Of these, 2,466,189 (74.4%) were successfully matched to H&E nuclear instances (**Fig. 2D-G**). Incomplete correspondence between the two sets of cellular instances may reflect differences in segmentation, spatial registration, and the constraints imposed by one-to-one matching.

### XenoType combines nuclear segmentation with independent histological representations

XenoType separates nuclear segmentation from the image representations used for classification (**Fig. 1D**). A state-of-the-art cell segmentation model (CellViT-SAM-H^13^) identifies nuclear instances, while an independently selected pathology foundation model extracts spatial image features. Crop selection and coordinate transformations associate each nuclear bounding box with the corresponding feature-token grid, and overlapping tokens are averaged to obtain a cell-level embedding. A classification head assigns a category to each cell based on its embedding to generate spatial classification maps.

A preliminary model configuration search evaluated combinations of 10 feature-extraction configurations, four classification heads, and three loss functions, yielding results for 95 of 120 possible combinations (**Supplementary Figs. S1 and S2**). The combination of H-Optimus-1^37^ with a multilayer perceptron and cross-entropy loss achieved the highest reported validation micro-F1 of 0.785, compared with 0.713 for the native CellViT-SAM-H configuration using the same head and loss. These experiments used earlier annotations supplied with the 10x dataset and preceded construction of the final four-reference annotation resource. The selected H-Optimus-1-MLP configuration was subsequently trained using the final molecularly inferred labels and evaluated on the eight held-out sections under the four- and seven-class taxonomies.

### XenoType improves four-class agreement with molecular reference annotations

We evaluated the classification across eight held-out breast tissue sections using four PanNuke-aligned categories: neoplastic, epithelial, connective, and inflammatory cells. We tuned the retrained CellViT++ classification head using the same hyperparameter optimization protocol as for XenoType. XenoType achieved a mean macro-F1 of 0.703 ± 0.091 and micro-F1 of 0.792 ± 0.072, compared with 0.569 ± 0.047 and 0.682 ± 0.075 for CellViT++ retrained on the molecular annotations, and 0.455 ± 0.058 and 0.607 ± 0.116 for default CellViT++, respectively (**Fig. 3C**). Values represent the mean ± sample standard deviation across sections. Both retrained models achieved higher mean scores than default CellViT++, with XenoType achieving the highest mean performance under both averaging conventions (**Supplementary Fig. S3**). For XenoType versus retrained CellViT++, the unadjusted P value was 0.00781 for both macro-F1 and micro-F1.

**Figure 3.**
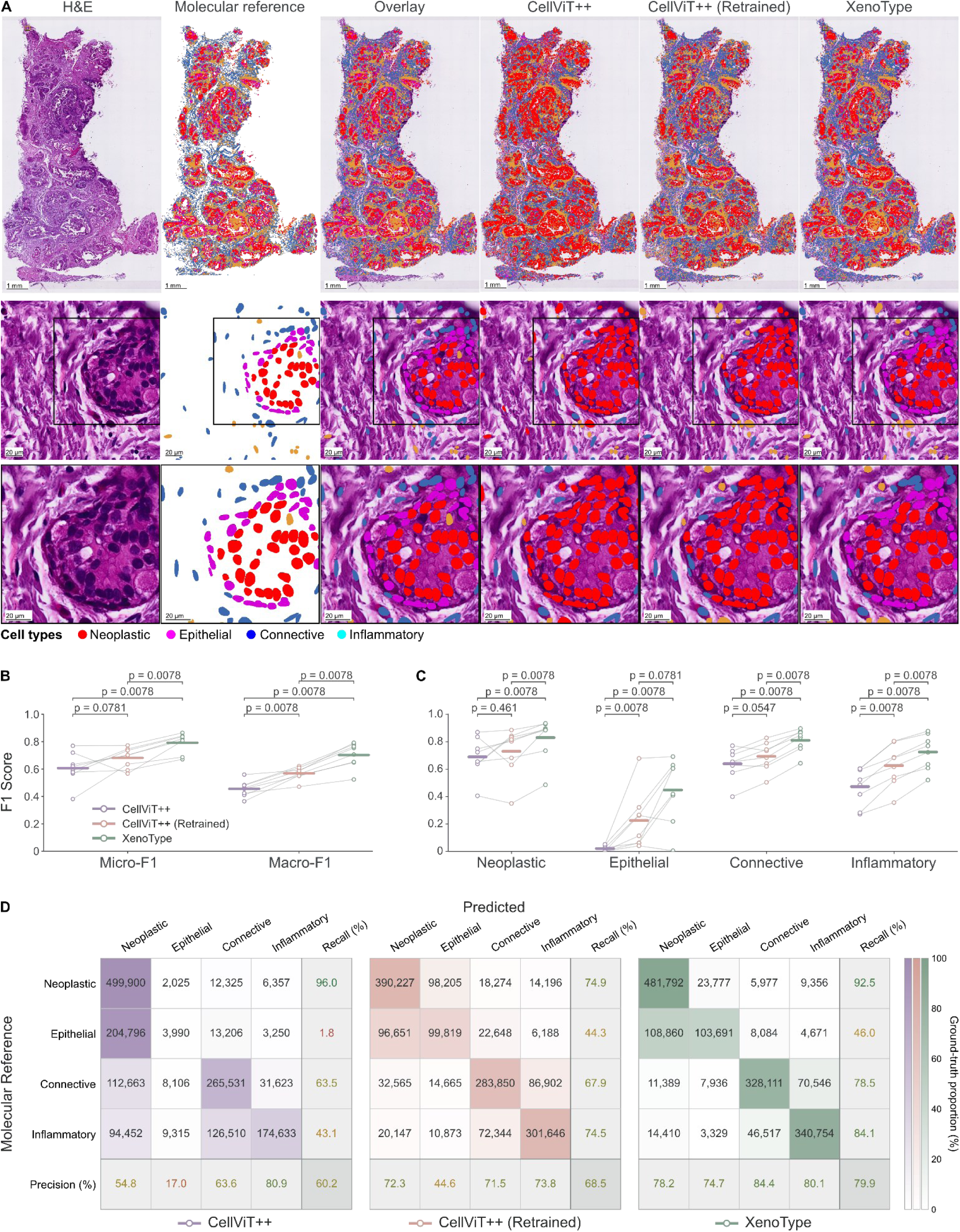
XenoType improves four-class agreement with molecular reference annotations across eight held-out breast tissue sections. **(A)** Whole-slide H&E image, molecular reference annotations, reference annotations overlaid on H&E, and predictions from default CellViT++, retrained CellViT++, and XenoType for an illustrative section (top row). Successive enlargements of the indicated region compare annotations and predictions at intermediate (middle row) and cellular (bottom row) scales across four PanNuke-aligned categories: neoplastic, epithelial, connective, and inflammatory cells. **(B)** Micro- and macro-averaged F1 scores across eight sections. (**C)** Class-specific F1 scores across the same sections. Bars indicate means, and points represent individual sections (n = 8). Brackets show unadjusted P values from exact, two-sided Wilcoxon signed-rank tests on scores paired by section (n = 8). **(D)** Confusion matrices pooled across the eight sections for default CellViT++, retrained CellViT++, and XenoType, from left to right. Rows indicate molecular reference categories and columns indicate predicted categories. Entries show cell counts, shading indicates row-normalized percentages, and marginal values report class-specific recall and precision.

XenoType achieved the highest mean class-specific F1 across all four categories (**Fig. 3C**). The largest increase over retrained CellViT++ occurred in the epithelial group, from 0.225 ± 0.213 to 0.447 ± 0.237, although this group remained the lowest-scoring category and showed substantial variation across sections. Mean F1 scores for neoplastic, connective, and inflammatory cells reached 0.830, 0.810, and 0.725, respectively.

Pooled confusion matrices summarize classification disagreements across the eight sections (**Fig. 3D**), whereas whole-slide maps and enlarged regions illustrate the spatial distribution of predictions in one section (**Fig. 3A**). Together, these results show improved agreement with molecular reference annotations under the four-class taxonomy, while identifying epithelial classification as a persistent limitation.

### XenoType improves seven-class agreement across held-out breast sections

We evaluated classification across the same eight held-out sections using seven molecularly informed categories. XenoType achieved a mean macro-F1 of 0.571 ± 0.058 and micro-F1 of 0.708 ± 0.064, compared with 0.403 ± 0.037 and 0.575 ± 0.063, respectively, for CellViT++ retrained on the same molecular annotations (**Fig. 4C**). Values represent the mean ± sample standard deviation across sections. XenoType achieved higher macro-F1 and micro-F1 than the retrained comparator on all eight sections (**Supplementary Fig. S3**). For XenoType versus retrained CellViT++, the unadjusted P-value was 0.00781 for both macro-F1 and micro-F1.

**Figure 4.**
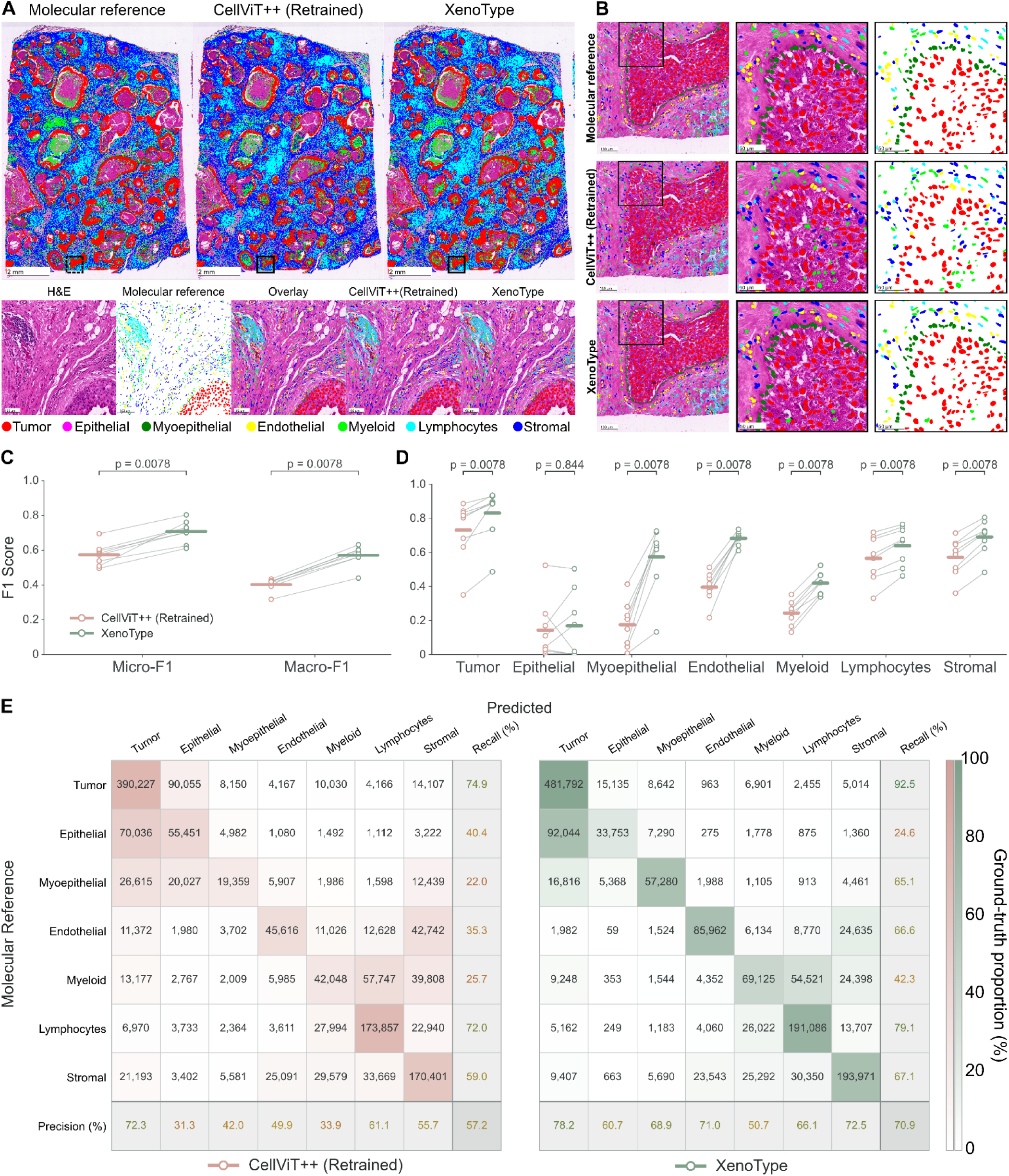
XenoType improves seven-class agreement with molecular reference annotations across eight held-out breast tissue sections. **(A)** Molecular referenceannotations and predictions from retrained CellViT++ and XenoType for an illustrative section (top row). An enlarged region (bottom row) compares the H&E image, molecular reference annotations, reference annotations overlaid on H&E, and model predictions across seven molecularly informed categories. **(B)** Further enlargement of the indicated region, showing molecular reference annotations, retrained CellViT++ predictions, and XenoType predictions as nuclear contours overlaid on H&E (left) and without the underlying image (right). Colors identify tumor, epithelial, myoepithelial, endothelial, myeloid, lymphocyte, and stromal cells. **(C)** Micro- and macro-averaged F1 scores for retrained CellViT++ and XenoType across eight sections. **(D)** Class-specific F1 scores across the same sections. In (C) and (D), points represent individual sections (n = 8), lines connect scores from the same section across models, and brackets show unadjusted P values from exact, two-sided Wilcoxon signed-rank tests on scores paired by section. **(E)** Confusion matrices pooled across the eight sections for retrained CellViT++ (left) and XenoType (right). Rows indicate molecular reference categories and columns indicate predicted categories. Entries show cell counts, shading indicates row-normalized percentages, and marginal values report class-specific recall and precision.

Mean F1 was higher for XenoType across all seven categories (**Fig. 4D**). The largest increases occurred for myoepithelial cells, from 0.175 to 0.572, and endothelial cells, from 0.395 to 0.682. Epithelial cells remained the lowest-scoring category, with mean F1 increasing from 0.142 ± 0.174 to 0.168 ± 0.198.

The pooled confusion matrices showed persistent disagreements involving epithelial and tumor cells, endothelial and stromal cells, and myeloid cells and lymphocytes (**Fig. 4E**). Whole-slide maps and enlargements illustrate the spatial distribution of reference annotations and predictions in one section (**Fig. 4A**).

### Classification agreement increases with annotation confidence and extends to ATERA-profiled breast tissue

Aggregate F1 scores weight all molecular reference annotations equally, regardless of the confidence with which they were assigned and, therefore, the reliability of the underlying label in the molecular reference annotations we construct. To investigate whether annotation confidence influences apparent model performance, we stratified results across the eight held-out sections by three proxies of annotation quality: the mean SingleR confidence score of the winning category across the four references used, the per-cell transcript count, and the number of uniquely detected genes (**Fig. 5A**). Micro-F1 increased consistently with all three measures, rising from approximately 0.40 at the lowest SingleR confidence scores to above 0.90 at the highest, with concordant trends for transcript counts and gene counts. The consistency of these relationships across three independent axes suggests that a substantial portion of model-reference disagreements concentrate on cells whose molecular annotations are themselves least certain. The relationship between annotation quality and classification accuracy varied across cell types (**Supplementary Fig. S7**).

**Figure 5.**
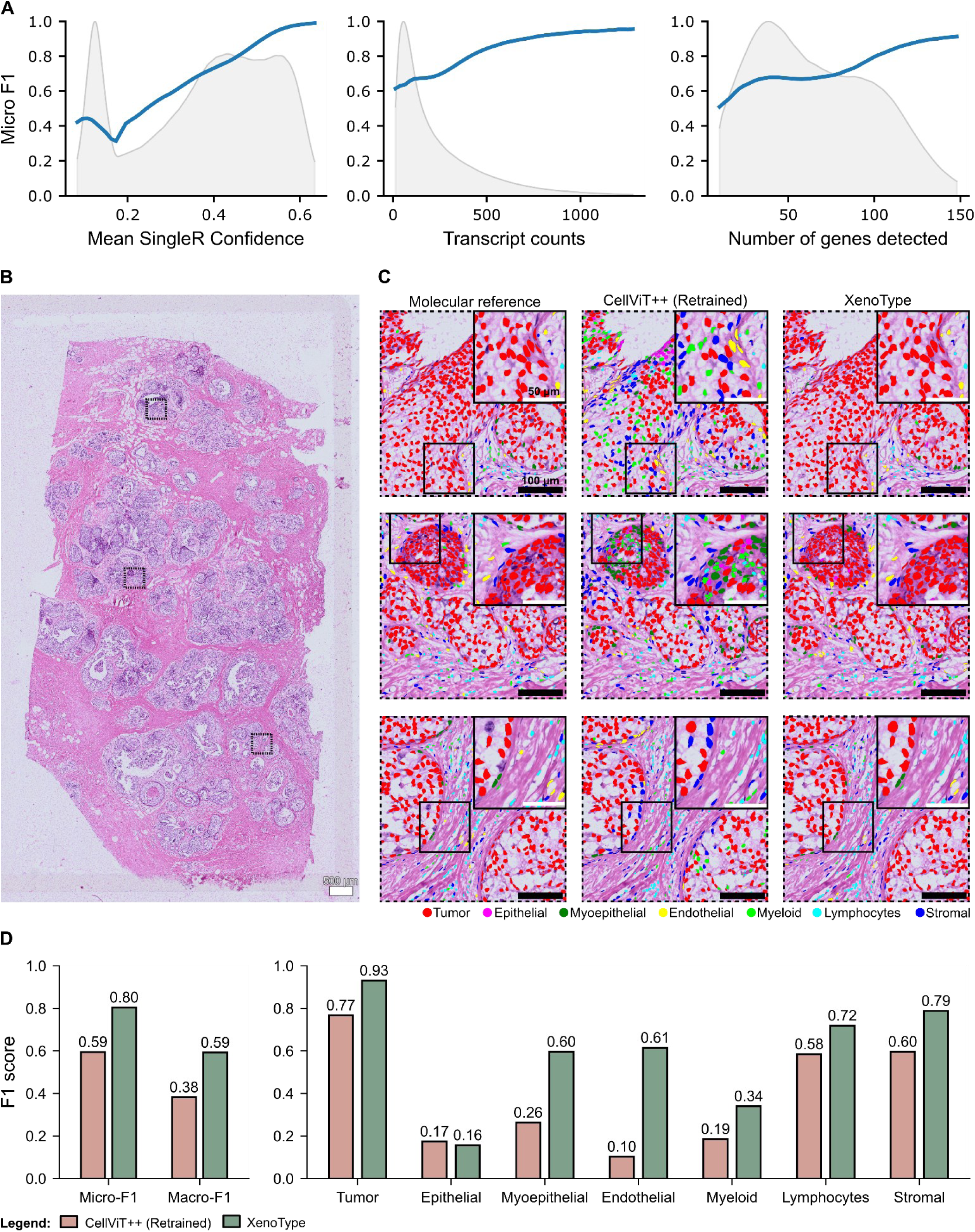
XenoType agreement with molecular references increases with annotation confidence and generalizes to out-of-distribution data. **(A)** Micro-F1 (blue curves) as a function of mean SingleR confidence, Xenium transcript count per cell, and number of detected genes per cell across the eight held-out breast tissue sections. Gray shading shows the distribution of each variable. **(B)** Whole-slide H&E image of the ATERA section. Dashed boxes mark the three regions shown in (C), ordered from top to bottom. **(C)** ATERA-derived molecular reference annotations (left), predictions from retrained CellViT++ (middle), and XenoType predictions (right), overlaid on the corresponding H&E regions. Colors denote the seven cell categories, and insets magnify the boxed subregions. Scale bars are 500 µm for the whole slide, 100 µm for the regional views, and 50 µm for the insets. **(D)** Micro-F1, macro-F1, and class-specific F1 scores on ATERA. XenoType achieved higher micro-F1 (0.80 vs. 0.59) and macro-F1 (0.59 vs. 0.38), with the largest gains for endothelial and myoepithelial cells.

Tumor, endothelial, and stromal cells showed strong positive trends across all three axes, with F1 exceeding 0.90 at the highest confidence scores, suggesting that low accuracy at the bottom of each axis may partly reflect uncertainty in molecular reference annotations rather than model failure. Lymphocyte and myeloid accuracy improved with annotation confidence but plateaued or declined at high transcript counts, consistent with atypical cell states or segmentation artifacts. Epithelial and myoepithelial cells showed inverted-U or declining patterns, indicating that the boundary between these populations and tumor cells remains ambiguous even for cells with the richest molecular profiles, which is consistent with the biological continuum between them and the reliance of both labels on a single reference.

To assess whether XenoType’s performance generalizes beyond the development cohort, we evaluated both XenoType and the retrained CellViT++ on an independent ATERA slide (**Fig. 5B**). ATERA is a whole-transcriptome, in situ spatial transcriptomics platform developed by 10x Genomics^38^. Cell-type annotations supplied with the dataset were harmonized into the seven cellular categories and used as molecular reference annotations. XenoType achieved a micro-F1 of 0.80 and macro-F1 of 0.59 on ATERA, compared with 0.59 and 0.38 for retrained CellViT++ (**Fig. 5D**). The largest class-specific gains were observed for endothelial and myoepithelial cells, consistent with the held-out evaluation. Epithelial cells remained the lowest-scoring category. Similar improvements held under the four-class taxonomy (**Supplementary Fig. S6**). Whole-slide spatial maps illustrated that XenoType predictions recapitulated tissue architecture more faithfully than retrained CellViT++ at both regional and cellular resolution (**Fig. 5B,C**).

## Discussion

Accurate identification of cellular populations in histological images is essential for characterizing the composition and spatial organization of the tumor microenvironment^3,39^. Deep learning and recent pathology foundation models have driven substantial progress in automated nuclear detection and classification^12,40^, yet cell classifiers remain constrained by training data that rely on labor-intensive manual labels, subject to observer variability and generally limited to broad categories such as neoplastic, epithelial, connective, and inflammatory cells^15,16^. Recent advances in imaging-based spatial transcriptomics and multiplex immunofluorescence now resolve molecular identities at single-cell resolution within intact tissue, creating new opportunities to supervise histological cell classification without manual labels^22,41,42^. Initial studies have demonstrated this approach using multiplex immunofluorescence-derived labels, while Xenium-based efforts such as STHELAR, STimage, and SHEST, have linked transcriptomic cell identities with corresponding H&E images for dataset construction and classifier development^18,23,24^. However, important limitations persist. Many spatial molecular assays either require a separate tissue section or alter the H&E stain itself, since H&E is typically performed after the molecular profiling rounds, producing images that differ from routine histological preparations and complicating the direct transfer of molecularly derived annotations to standard clinical-grade H&E. At the same time, leading deep learning models for histological image analysis have been developed primarily for nuclear segmentation and derive their classification features from the same segmentation backbone, limiting the independent selection of representations best suited for distinguishing finer cellular populations^12,13,32^. These constraints have kept histological cell classification largely confined to broad morphological categories.

Here, we developed XenoType, a framework to classify cells in routine H&E images using training annotations derived from same-section Xenium spatial transcriptomics. Cell-type labels were reconciled from four independent scRNA-seq references using SingleR, consistent with a previous benchmark identifying it as the best-performing method among five evaluated approaches on Xenium data^28^. Decoupling feature extraction from segmentation allowed the independent selection of a pathology foundation model for cell-level representations, addressing a limitation of segmentation-centric architectures. XenoType outperformed CellViT++ retrained on the same labels under both the four-class and seven-class taxonomies, demonstrating that molecularly supervised classification can extend beyond conventional broad categories to finer cellular populations. XenoType’s improved performance over retrained CellViT++ on the ATERA section, using cell-type annotations generated independently of our multi-reference annotation procedure, suggests that its performance gains are not specific to the annotation strategy used for training. Because molecular measurements are required only during dataset construction, inference uses the H&E image alone, supporting scalable cell-type mapping without per-specimen molecular profiling. The same-section spatial transcriptomics design reduces registration uncertainties and staining artifacts inherent to serial-section or post-IF H&E workflows, enabling direct annotation transfer to standard-quality histological images.

Several limitations should be noted. First, epithelial cells remained the worst performing category, and per-class analysis confirmed that accuracy did not improve with increasing transcriptomic depth (**Supplementary Fig. S7**), suggesting that uncertainty in the reference annotations may contribute to the observed classification errors. The ∼280-gene Xenium panel may be insufficient to resolve epithelial from tumor cells - a distinction that also depends on copy number alterations and somatic mutations not captured by targeted expression panels^43^. Furthermore, off-target probe binding has been documented in current Xenium breast cancer gene panels, potentially introducing additional uncertainty into transcriptomics-derived cell annotations^44^. In addition, only one reference (Wu *et al.* ^2^) contained epithelial and myoepithelial categories, leaving these labels without the multi-reference support available for other populations^2,45^, consistent with the trends observed for both classes (**Supplementary Fig. S7**). More broadly, model-reference agreement increased consistently with annotation confidence, although class-specific trends varied (**Supplementary Fig. S7**), indicating that aggregate F1 scores partly reflect ground-truth uncertainty rather than model limitations alone. Second, the development and held-out evaluation data originated from a single public collection. Although evaluation sections came from different patients than the development sections, shared sample preparation, staining, and imaging conditions limit the independence of the comparison. The ATERA slide, profiled on a different spatial transcriptomic platform, provides an initial out-of-distribution test, but multi-institutional evaluation with diverse tissue preparation and staining protocols remains needed. Finally, XenoType and retrained CellViT++ differ simultaneously in feature extractor and tiling strategy, so the observed improvements cannot be attributed to any single component without controlled ablation experiments^13^.

A number of directions could extend this work. Although the ATERA section provides initial cross-platform evidence, evaluation on external cohorts from independent institutions remains needed to establish generalizability. Extending the framework to other tissue types - such as colorectal, lung, or pancreatic cancers - would test whether molecularly supervised classification transfers across different tumor microenvironments, while larger training datasets aggregating multiple spatial transcriptomics platforms could improve robustness. A key goal is to develop even finer cell categories, such as cancer molecular subtypes or functional immune states. The annotation-confidence analysis (**Fig. 5A**) indicates that richer molecular profiles are associated with greater model-reference agreement, suggesting broader gene panels, integration of copy number variation and mutational information, and independent protein validation will be essential for this goal. Finally, coupling spatially resolved cell-type predictions with clinical metadata could link cellular composition to patient survival and treatment response, moving from methodological validation toward biological and clinical discovery.

In summary, XenoType demonstrates that spatial transcriptomics-derived annotations can supervise histological classifiers capable of distinguishing molecularly informed cellular populations beyond conventional morphological categories. Model-reference agreement was correlated with annotation confidence, indicating that aggregate F1 scores partly reflect ground-truth uncertainty, and initial out-of-distribution evaluation supported cross-platform generalization. Although broader validation remains necessary, these findings support the potential of molecularly supervised learning to extend the cellular resolution of conventional histology and provide a foundation for investigating tissue heterogeneity at scale.

## Methods

### Data acquisition and image preprocessing

We used the publicly available Xenium v1 Human Breast FFPE with Biomarkers & Housekeeping Genes Custom Panel dataset generated by 10x Genomics^46^. The collection comprises 12 human breast tissue sections from 12 donors, including eight ductal carcinoma *in situ* (DCIS) specimens, three invasive ductal carcinomas (IDC), and one normal breast specimen. Tissue sections were profiled using Xenium In Situ with a custom 280-gene panel and subsequently stained and imaged with H&E, providing paired spatial transcriptomic and histological data. Publicly available cell-level expression matrices, transcript coordinates, nuclear boundaries, and H&E whole-slide images were used for downstream analysis. Specimen characteristics and acquisition procedures are described in the original dataset publication^19^.

Spatial correspondence between Xenium measurements and H&E images was established through manual landmark-based registration using Xenium Explorer (10x Genomics). Corresponding anatomical landmarks were identified in the Xenium DAPI and H&E images, from which slide-specific similarity transformations were estimated by least-squares fitting. The resulting 3 x 3 affine transformation matrices were exported and used, together with the corresponding image pixel resolutions, to map cell and nuclear coordinates between the two imaging modalities. Four tissue sections were selected for model development, while the remaining eight sections were reserved for independent evaluation and excluded from training, validation, and hyperparameter selection. Section identifiers, specimen characteristics, and experimental allocation are provided in **Supplementary Table S1**.

For external evaluation, we used a publicly available FFPE breast cancer section profiled using the preproduction ATERA Whole Transcriptome Assay (10x Genomics)^47^. The specimen comprised ductal carcinoma in situ (ER+/PR+/HER2-), with spatial expression measurements targeting 18,028 genes and corresponding H&E imaging. The section was not used for model development or hyperparameter selection.

### Single-cell reference preparation

Four breast cancer single-cell RNA-sequencing (scRNA-seq) reference datasets (Wu^2^, Chen^33^, Dong^34^, and Yan^35^) were used for molecular cell-type annotation. Cell identities were obtained from the original annotations provided with each reference dataset and harmonized into seven cellular categories: tumor, epithelial, myoepithelial, endothelial, myeloid, lymphocyte, and stromal cells. Reference accessions, original cell-type annotations, and their correspondence to the harmonized categories are provided in **Supplementary Table S2**.

Reference cells were retained if they contained at least 200 detected genes and 500 counts, with mitochondrial counts comprising no more than 20% of the total. Cells with missing or uninformative annotations were excluded. Expression data were normalized using log-normalization (scale factor = 10,000), and each reference was restricted to genes shared with the Xenium panel. Reference sampling was capped at 15,000 cells per label. Principal component analysis and UMAP were used to visualize reference structure.

### Xenium cell annotation and reference agreement

We retained Xenium cells with at least 10 transcripts and 5 detected genes. We annotated each eligible expression profile independently against the four single-cell RNA-sequencing references using SingleR^36^. We mapped the reference-specific predictions to the seven-category taxonomy and combined them by voting. We resolved ties using the mean SingleR score for each tied category, followed by the predefined reference priority order (Yan, Wu, Dong, and Chen) if a tie remained. When the Wu reference assigned a cell to the epithelial or myoepithelial category, we used that assignment as the final label.

We counted the number of references that supported each final assignment and classified the agreement into high (four reference agreement), medium (three reference agreement), and low (fewer than three reference agreement). Because each reference could support only the categories represented in its mapped labels, the maximum possible agreement varied by category. **Supplementary Table S2** lists the categories available in each reference.

### Nuclear segmentation and annotation transfer

Nuclear instances were detected in H&E whole-slide images using the pretrained CellViT-SAM-H-x40-AMP model^13^. To associate histologically detected nuclei with molecularly annotated Xenium cells, spatial correspondence was established using slide-specific affine transformation matrices. Nuclear coordinates were transformed into the Xenium coordinate system for nearest-centroid matching, while the resulting annotations were retained in the H&E coordinate system.

Each detected nucleus was matched to its nearest Xenium cell using a k-dimensional tree, with a maximum matching distance of 15 µm. When multiple detected nuclei were associated with the same Xenium cell, only the closest match was retained. A 10-µm margin around the Xenium field of view was applied during spatial eligibility filtering. Successfully matched H&E nuclei inherited the corresponding Xenium cell identifiers and molecularly inferred cell-type labels, while the original segmentation was preserved for subsequent classifier training and evaluation.

### Tile extraction and dataset curation

Candidate image tiles measuring 1,024 x 1,024 pixels were extracted at a stride of 256 pixels. Tiles containing fewer than 10 annotated cells or more than 30% whitespace were excluded, with whitespace filtering performed using an intensity threshold of 220 (**Supplementary Fig. S4**). An additional nuclear area filter was configured to identify unusually large instances using a threshold of three standard deviations above the mean. Candidate selection incorporated Shannon entropy calculated from the proportions of annotated cell categories, favoring tiles containing diverse cellular populations.

A greedy selection procedure was used to construct spatially nonoverlapping tile subsets, with an eight-pixel separation buffer. The resulting dataset comprised 372 training tiles and 66 validation tiles, containing 122,398 and 19,935 annotated cells, respectively, according to the dataset creation report. Sampling targets of approximately 21,000 training cells and 4,300 validation cells per category guided tile selection, although the final class distributions remained unequal. All seven categories were represented in both partitions. Training and validation tiles originated from the same four specimens but occupied nonoverlapping image regions. The final class distributions and slide-level allocations are reported in **Supplementary Table S3**.

### XenoType architecture and cellular feature assignment

XenoType decouples nuclear segmentation from histological feature extraction by combining CellViT-SAM-H-derived nuclear instances with representations generated by an independently selected pathology foundation model. The selected configuration used H-Optimus-1 as a frozen feature extractor and a multilayer perceptron (MLP) for seven-class cell classification^48^.

For H-Optimus-1 feature extraction, CellViT-normalized H&E images were restored to RGB intensity values and resampled to 0.5 µm/pixel using bilinear interpolation. The rescaled images were divided into overlapping 224 by 224-pixel crops, with crop positions distributed uniformly to ensure complete image coverage without padding and a maximum stride of 112 pixels and H-Optimus-1 was applied independently to each crop. Auxiliary transformer tokens were excluded, and the remaining spatial tokens were organized into a 16 x 16 grid of 1,536-dimensional representations.

To associate these representations with individual nuclei, CellViT-SAM-H bounding boxes and centroids were transformed into the rescaled image coordinate system. Each nucleus was preferentially assigned to a 224 x 224-pixel H&E crop (112 x 112 µm at 0.5 µm/pixel) that fully contained its bounding box. If multiple crops qualified, we selected the one with the greatest minimum distance between the bounding box and the crop edges. When no crop fully contained the bounding box, the crop containing the nuclear centroid with the greatest boundary margin was selected. The bounding box was subsequently mapped onto the spatial token grid, and the embeddings of all tokens whose image patches intersected the bounding box were averaged to generate a single 1,536-dimensional embedding per nucleus. Cell ordering was preserved throughout feature extraction to maintain correspondence with the original nuclear detections.

The resulting embeddings were passed to an MLP comprising a 64-unit hidden layer, rectified linear activation, dropout, and a seven-class output layer. Predicted cell identities were associated with their corresponding nuclear instances to generate spatially resolved classification maps.

### Model training and configuration selection

To select the architecture described above, we first compared combinations of 10 feature-extraction configurations , four classification heads, and three loss functions, completing 95 of the 120 possible combinations (**Supplementary Figs. S1 and S2**). This preliminary search used the original Xenium cell-type annotations harmonized into the seven-category taxonomy (**Supplementary Fig. S5**). H-Optimus-1 with an MLP head and cross-entropy loss achieved the highest validation micro-F1 (0.785) among the evaluated combinations and was selected for subsequent training. Hyperparameter optimization and experiment tracking were performed using Weights & Biases^49^.

We then trained the selected H-Optimus-1-MLP configuration on the final SingleR-derived labels for seven-class classification using cross-entropy loss and AdamW optimizer. The classifier used a 64-unit hidden layer with a dropout probability of 0.4544. Training was configured for a maximum of 100 epochs, with a batch size of 256, learning rate of 3.012 × 10⁻⁴, weight decay of 0.0253, and AdamW momentum parameters β₁ = 0.9 and β₂ = 0.999. Cosine learning-rate scheduling, mixed-precision computation, and early stopping with a patience of five validation epochs were applied. A random seed of 15 was used, and class-weighted sampling was disabled. Histological feature vectors were cached, and the classification head was optimized using the resulting cell-level embeddings.

The checkpoint with the highest validation micro-F1 score was retained for evaluation. The best checkpoint was obtained at epoch 19, with a validation F1 of 0.7545, and training stopped after epoch 24.

### Model evaluation

XenoType was compared with default CellViT++ and a CellViT++ classifier retrained using Xenium-derived molecular annotations. Classification was evaluated using four broad cellular categories and the seven molecularly informed categories. For the four-class evaluation, tumor cells were assigned to the neoplastic category, epithelial and myoepithelial cells to epithelial, endothelial and stromal cells to connective, and myeloid cells and lymphocytes to inflammatory.

For each of the eight held-out sections, we matched predicted nuclei to H&E nuclear instances carrying transferred molecular annotations. We identified candidate pairs with centroid distances of at most 15 pixels in the H&E coordinate system and accepted pairs in ascending order of distance, allowing each instance to be matched at most once. We performed matching separately for each model and computed classification scores on its matched pairs. In the four-class evaluation, we excluded pairs with predictions labeled Dead (from CellViT++) after matching.

We computed micro-F1, macro-F1, and class-specific F1 for each section using the TorchMetrics multiclass F1 implementation. We report the unweighted mean and sample standard deviation of the section-level scores and pooled matched-cell counts across sections before calculating row-normalized percentages.

We compared models using exact, two-sided Wilcoxon signed-rank tests on F1 scores paired by tissue section (eight sections). The four-class analysis included all three model pairs, and the seven-class analysis compared XenoType with retrained CellViT++. For each model pair, we tested micro-F1, macro-F1, and class-specific F1. We excluded zero paired differences, assigned average ranks to tied absolute differences, and enumerated all sign assignments to calculate exact P values. We report unadjusted P values and interpret the comparisons as exploratory.

To assess cross-platform generalization, the models were additionally evaluated on the ATERA breast cancer section without further training. Cell-type annotations provided by 10x Genomics were harmonized into the seven cellular categories and transferred to H&E nuclear instances. XenoType and retrained CellViT++ were evaluated under both the four- and seven-class taxonomies, with default CellViT++ additionally included in the four-class comparison. Classification performance was quantified using the same centroid-matching procedure and F1 metrics described above.

### Annotation-confidence analysis

To examine the relationship between annotation confidence and classification performance, we stratified matched cells from the eight held-out sections by three proxies of annotation quality: the mean SingleR confidence score of the winning category across the four references, the per-cell Xenium transcript count, and the number of uniquely detected genes. Per-cell transcript counts and gene counts were obtained from the Xenium cell-level expression data. For each axis, values were clipped to the 1st-99th percentile range and evaluated at 50 evenly spaced thresholds using a sliding window with a half-width of 5% of the clipped range. Windows containing fewer than 1,000 cells were excluded. Micro-F1 and macro-F1 were computed within each window, and uncertainty was estimated by bootstrap resampling (50 iterations, seed = 42). The background distribution of each variable was estimated using Gaussian kernel density estimation. The analysis was performed on all seven classes. For the per-class analysis (**Supplementary Fig. S7**), class-specific F1 score was computed using the same sliding-window procedure with a minimum of 200 cells per window, and bootstrap standard deviations were used to generate ±1 SD bands.

### Software and data availability

Spatial transcriptomic processing, reference-based annotation, and histological image analysis were implemented using R and Python. SingleR was used for reference-based cell annotation, while PyTorch-based implementations of CellViT-SAM-H, H-Optimus-1, and the classification heads were used for histological inference and model training. Training configurations, metrics, and experiment metadata were recorded using Weights & Biases.

The Xenium v1 Human Breast FFPE with Biomarkers & Housekeeping Genes Custom Panel dataset^46^ and the ATERA In Situ Gene Expression FFPE Human Breast Cancer preview dataset^38^, together with their paired H&E images, are publicly available from 10x Genomics. The XenoType implementation, trained checkpoints, derived molecular annotations, and processed histological datasets are available upon request.

## Supplementary Figures

**Supplementary Fig. S1.**
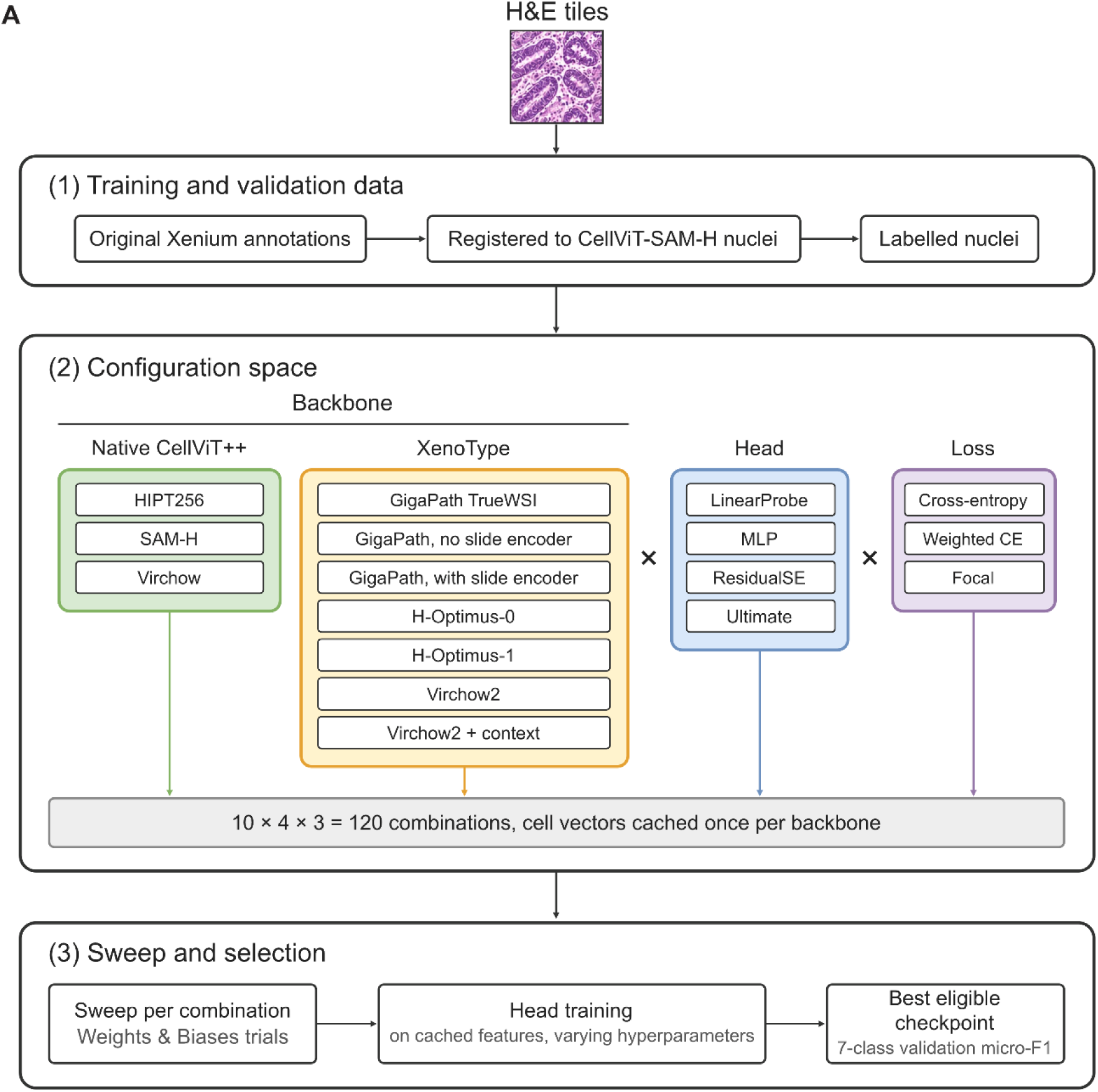
Preliminary configuration search for feature extractor, classifier head, and loss function selection. **(A)** Overview of the configuration search pipeline. (1) Training and validation data were constructed by registering original Xenium cell-type annotations to CellViT-SAM-H nuclear instances to produce labeled nuclei. (2) The configuration space comprised 10 feature-extraction backbones (3 native CellViT++ and 7 XenoType-compatible pathology foundation models), 4 classification heads (LinearProbe, MLP, ResidualSE, and Ultimate), and 3 loss functions (cross-entropy, weighted cross-entropy, and focal), yielding 120 possible combinations. Cell-level feature vectors were cached once per backbone. (3) For each combination, a hyperparameter sweep was conducted using Weights & Biases, with classifier heads trained on the cached features. The best eligible checkpoint was selected by seven-class validation micro-F1. This search used annotations supplied with the 10x dataset and preceded construction of the final four-reference annotation resource.

**Supplementary Fig. S2.**
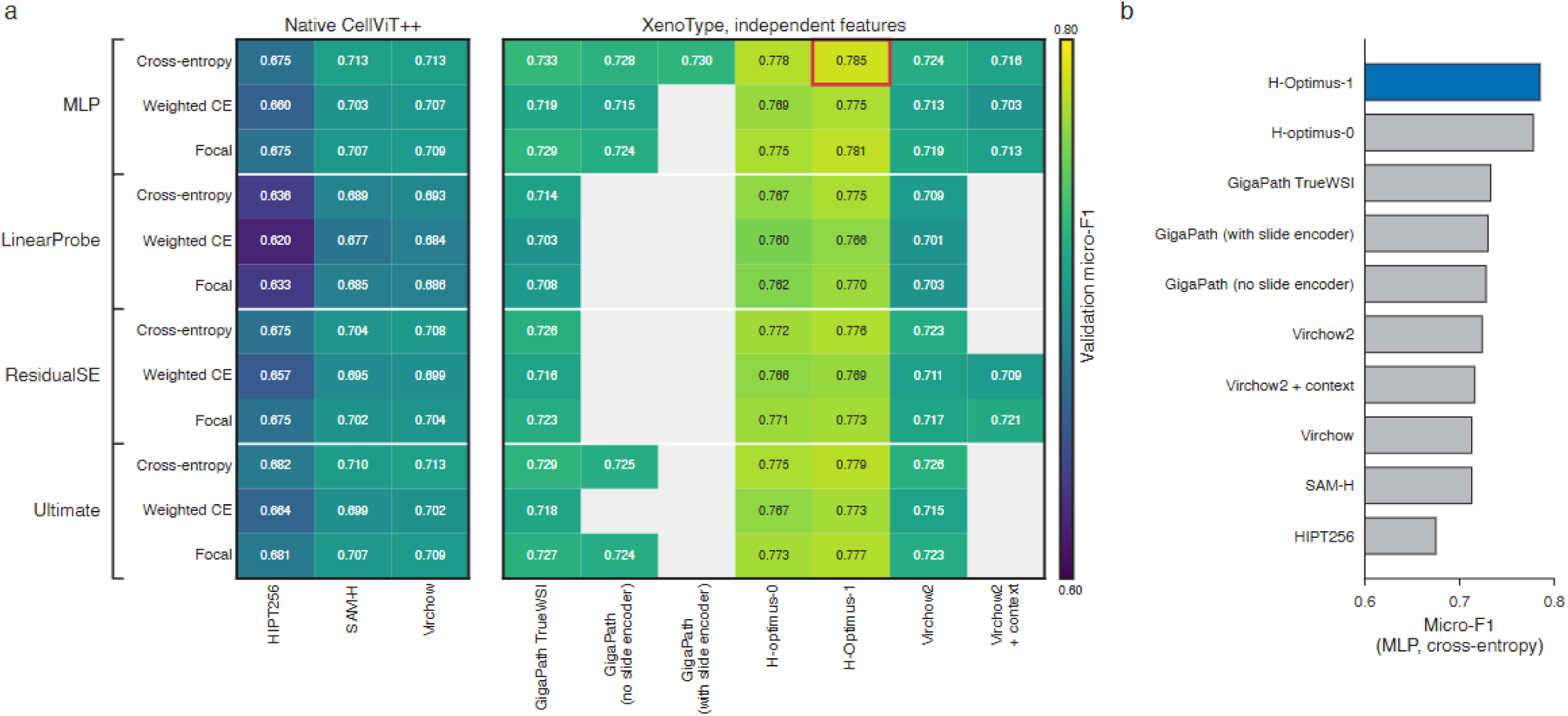
Validation performance in the preliminary configuration search. **(A)** Seven-class validation micro-F1 for 95 reported combinations of feature extractor, classifier head, and loss function. Gray cells indicate combinations with no reported result. **(B)** Comparison of the ten feature-extraction configurations using a multilayer perceptron and cross-entropy loss. The search used the earlier annotations supplied with the 10x dataset.

**Supplementary Fig. S3.**
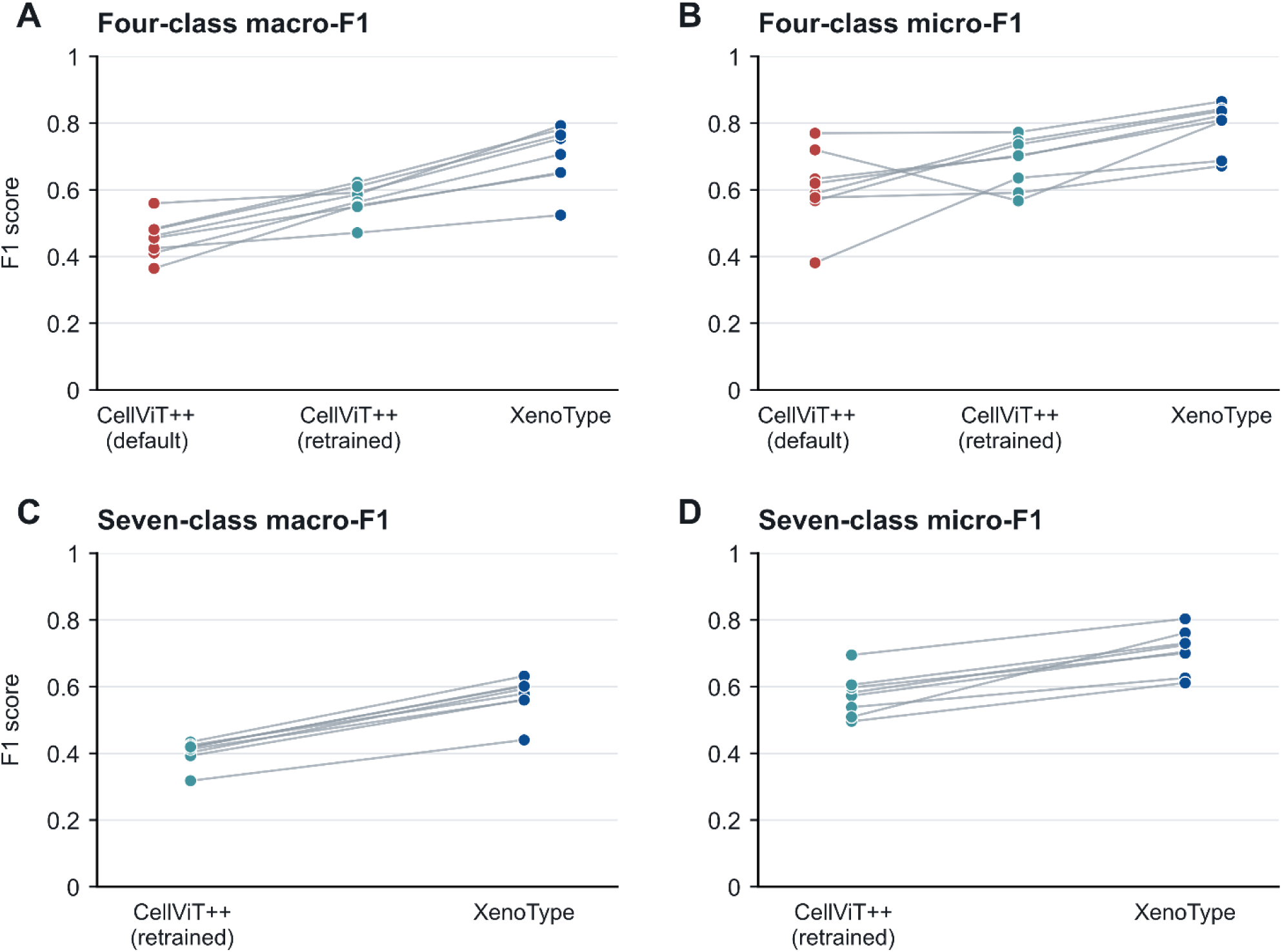
Paired model performance across eight held-out breast tissue sections. (A,B) Four-class macro-F1 and micro-F1 for default CellViT++, retrained CellViT++, and XenoType**. (C,D)** Seven-class macro-F1 and micro-F1 for retrained CellViT++ and XenoType. Each point represents one section, and lines connect scores from the same section across models.

**Supplementary Fig. S4.**
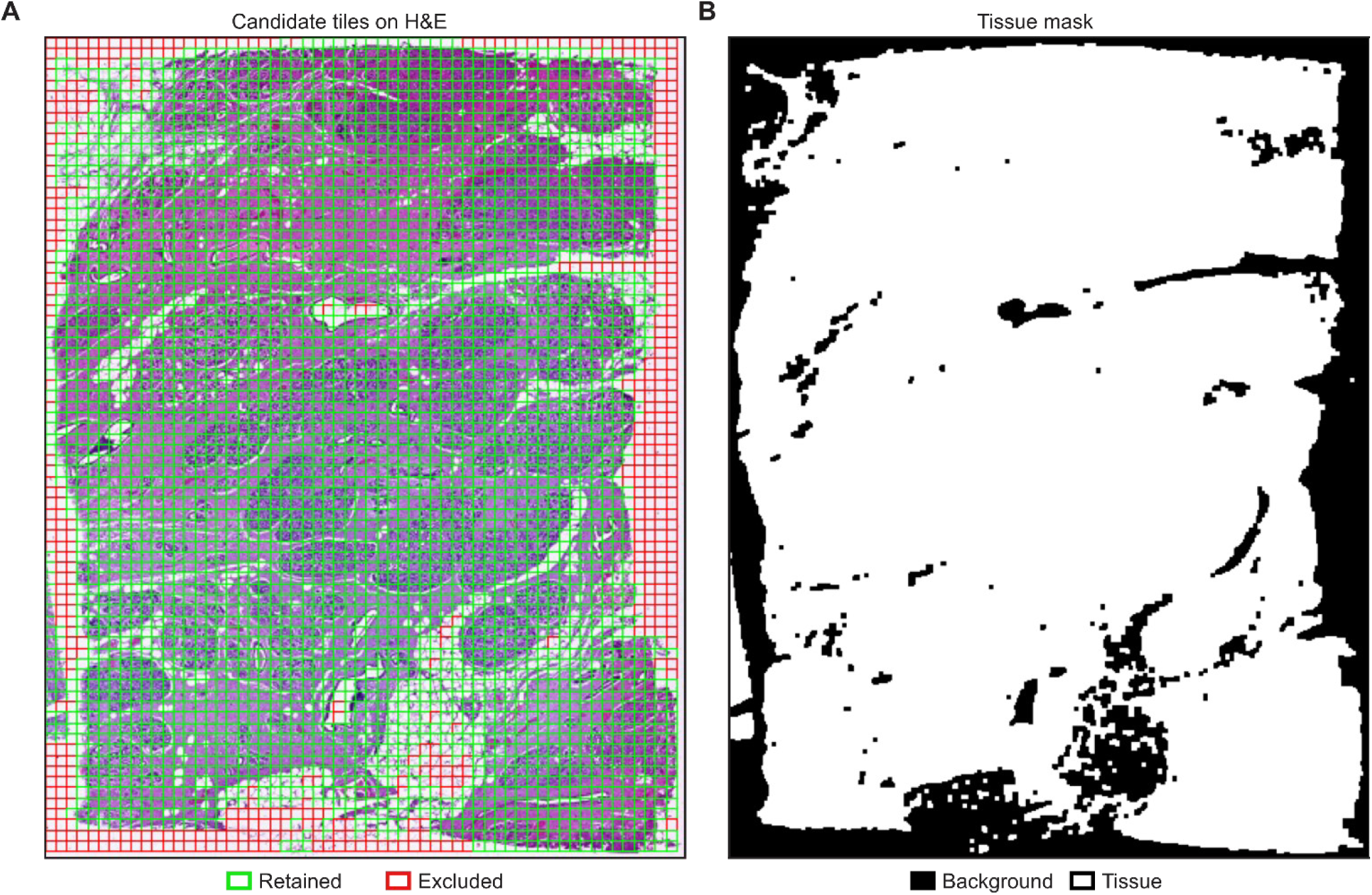
Tile extraction and whitespace filtering for a representative development section. **(A)** Candidate tiles overlaid on the H&E whole-slide image. Green and red borders indicate retained and excluded tiles, respectively. Tiles were excluded if they contained fewer than 10 annotated cells or more than 30% whitespace. **(B)** Binary tissue mask used for whitespace detection, generated using an intensity threshold of 220. White and black regions indicate tissue and background, respectively.

**Supplementary Fig. S5.**
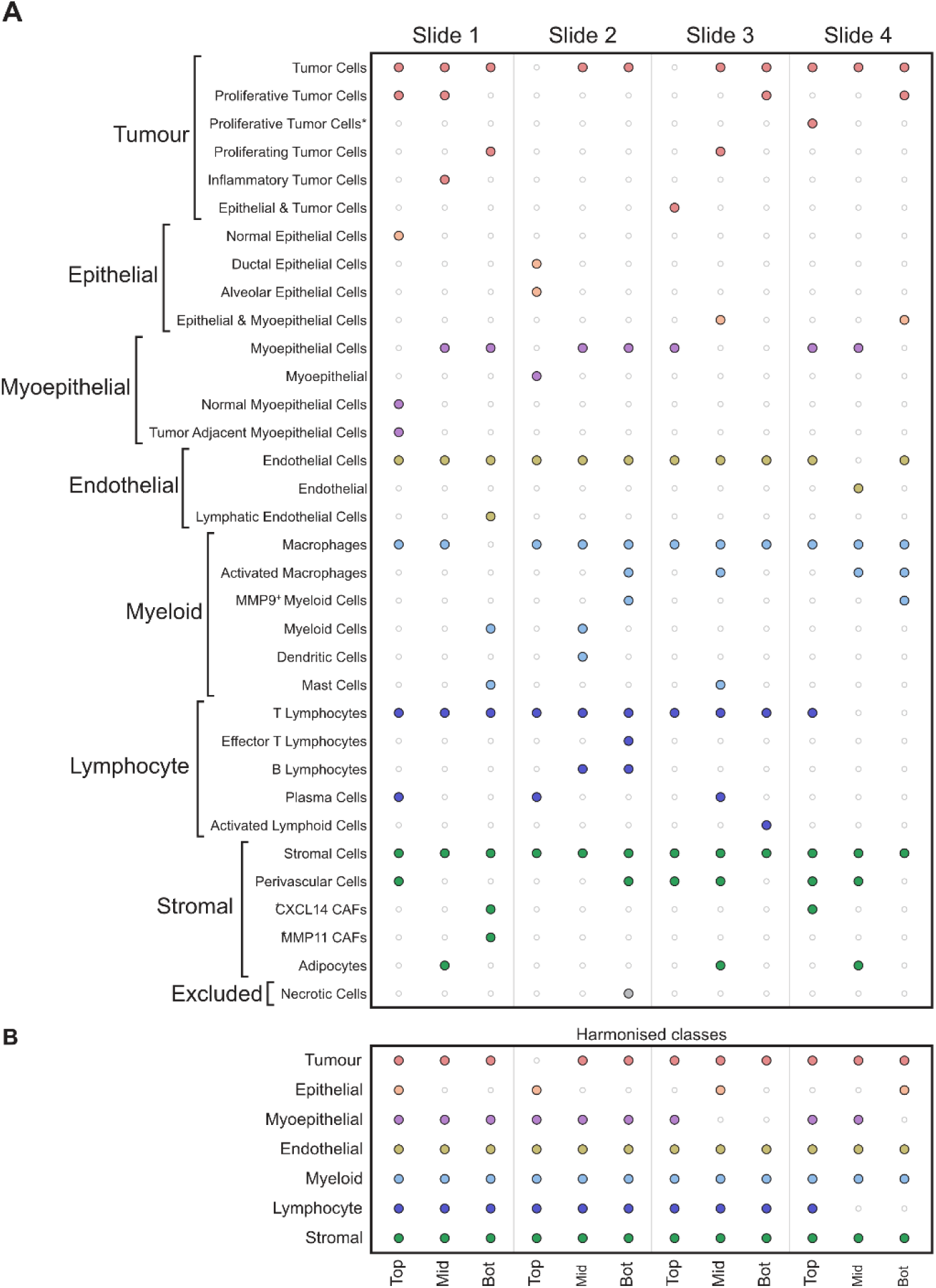
Original Xenium cell-type annotations and harmonized categories across 12 breast tissue sections. **(A)** Presence of fine-grained cell-type labels supplied with the 10x Genomics Xenium dataset across 12 tissue sections (four slides, three sections each). Labels are grouped by their corresponding harmonized category (left brackets). Dots indicate that a given label was assigned to cells in that section; colors correspond to the harmonized category. Necrotic cells (bottom) were excluded from downstream analysis. **(B)** Harmonized seven-category representation across the same sections, after collapsing the fine-grained labels shown in **(A)**.

**Supplementary Fig. S6.**
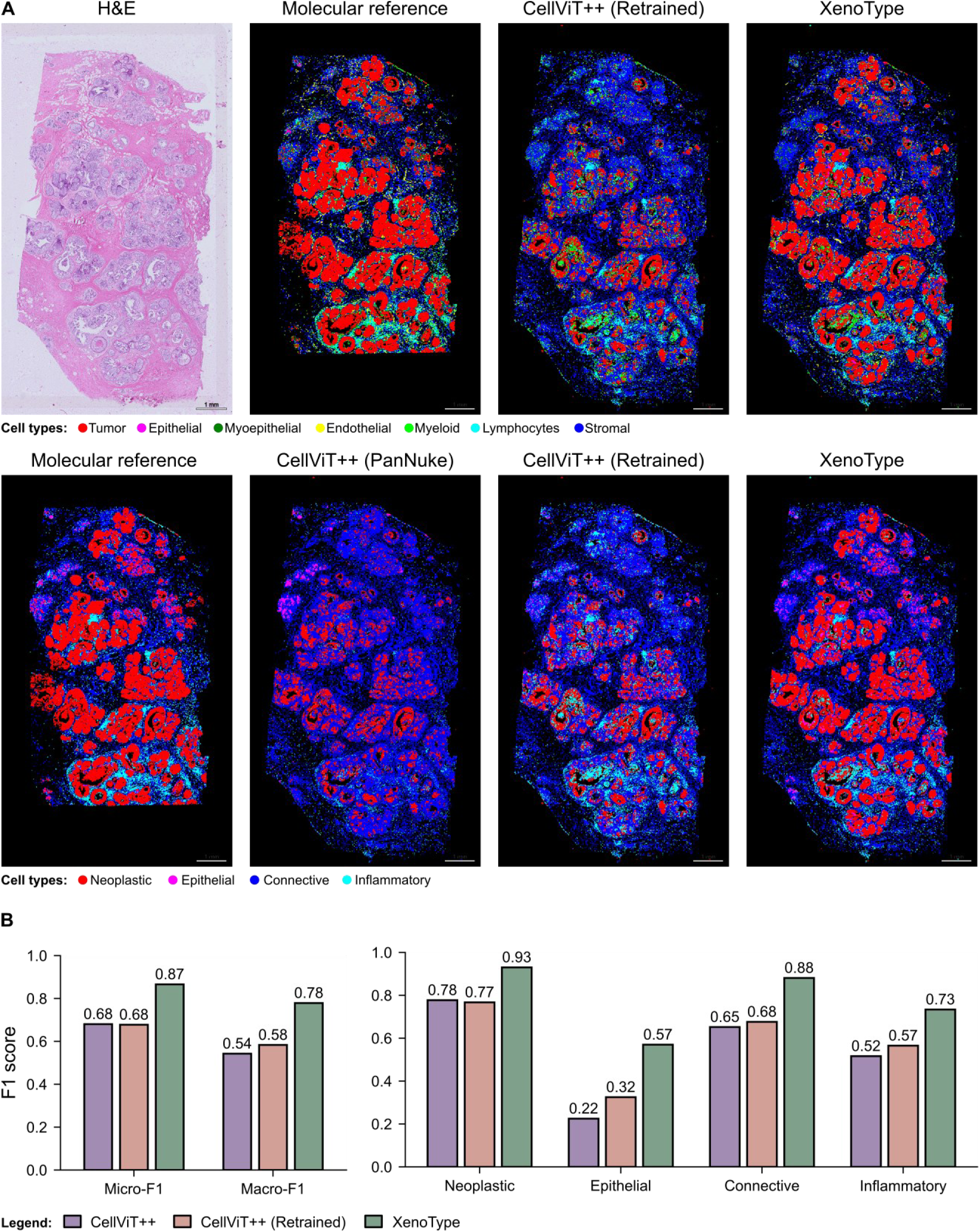
XenoType achieves higher F1 scores than both CellViT++ baselines across all four broad cell categories on ATERA. **(A)** Whole-slide H&E image and cell-type maps for the out-of-distribution ATERA breast tissue section. The top row shows the H&E image, ATERA-derived molecular reference annotations, and predictions from retrained CellViT++ and XenoType under the seven-class taxonomy. The bottom row shows the molecular reference annotations and predictions from PanNuke-trained CellViT++, retrained CellViT++, and XenoType mapped to four broad categories. Scale bars, 1 mm. **(B)** Micro-F1, macro-F1, and class-specific F1 scores against the molecular reference annotations under the four-class taxonomy.

**Supplementary Fig. S7.**
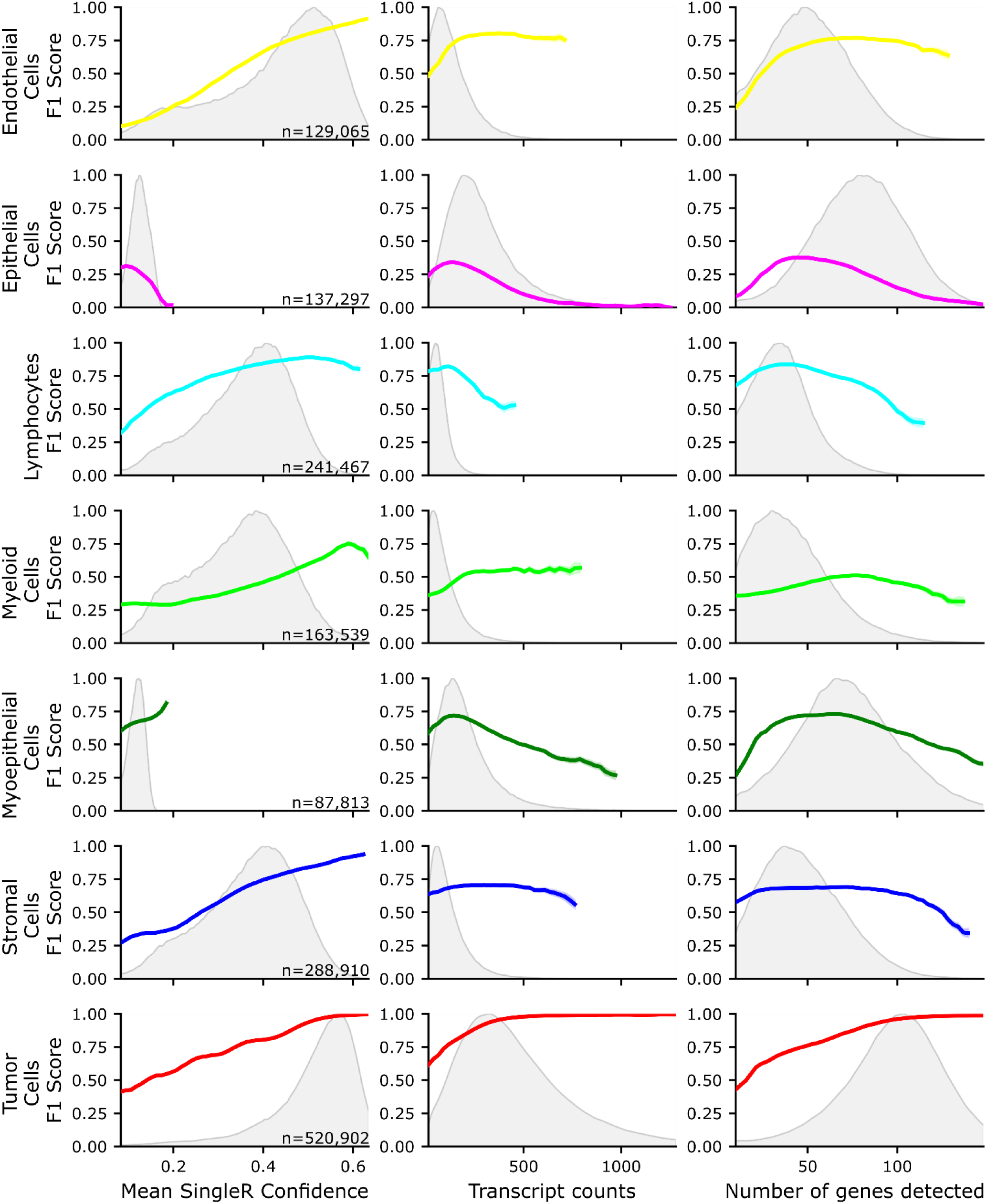
XenoType’s agreement with molecular reference annotations generally increases with SingleR confidence, while trends with transcript and gene counts vary by cell type. Class-specific F1 scores across the eight held-out breast tissue sections are shown as a function of mean SingleR confidence (left), Xenium transcript count per cell (middle), and number of detected genes per cell (right). Colored curves show F1 scores, and gray shading shows the distribution of each variable within the corresponding molecular reference category. Row labels give the number of matched nuclei per reference category.

## Supplementary Tables

**Table S1.**
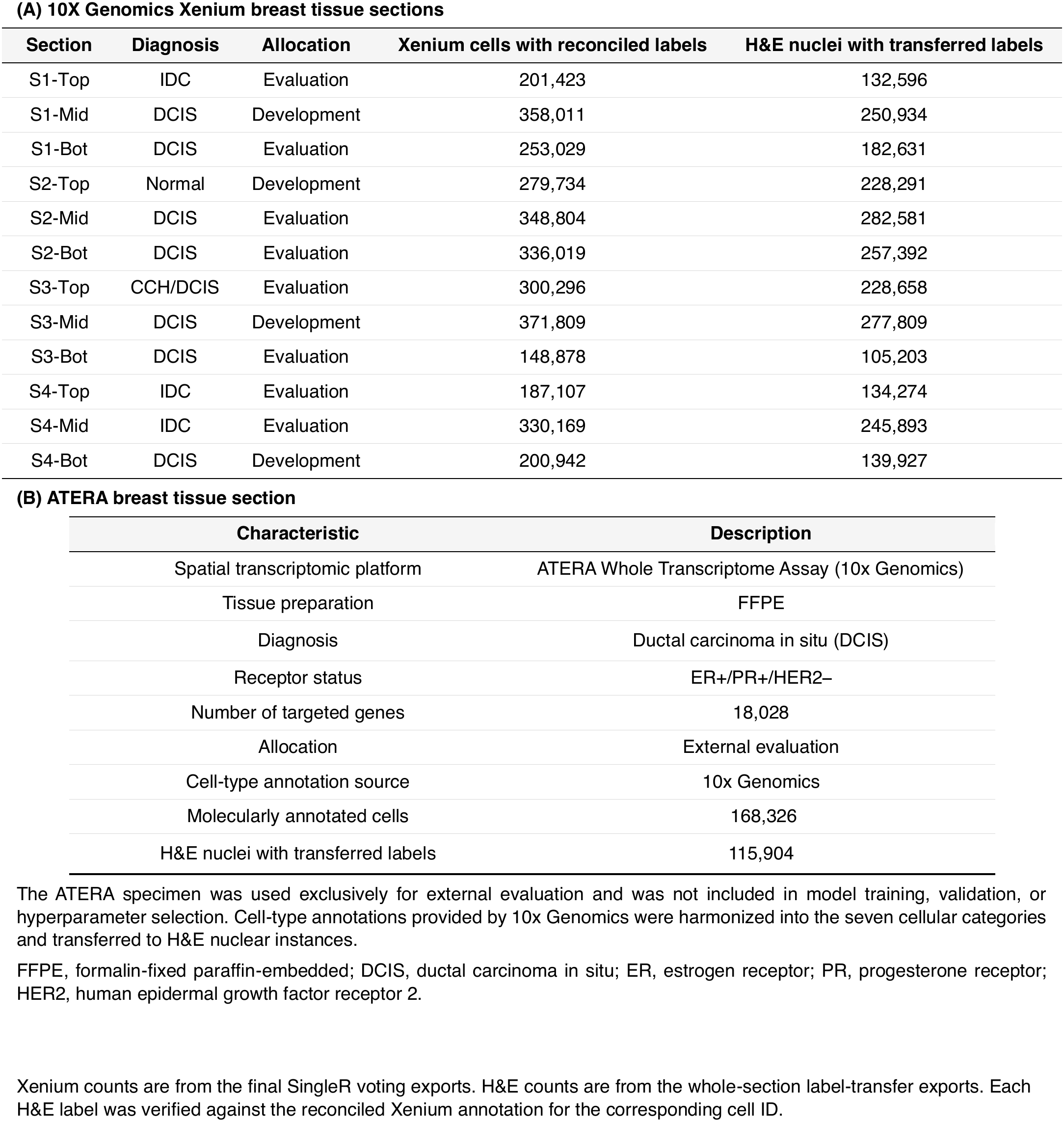
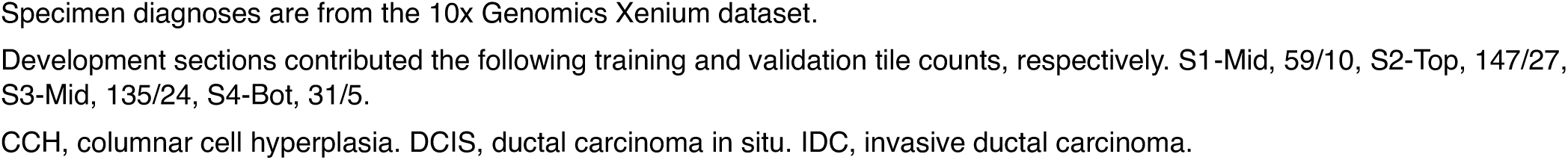
Breast tissue specimens and study allocation. Related to Figure 1.

| (A) 10X Genomics Xenium breast tissue sections |  |  |  |  |
| Section | Diagnosis | Allocation | Xenium cells with reconciled labels | H&E nuclei with transferred labels |
|---|---|---|---|---|
| S1-Top | IDC | Evaluation | 201,423 | 132,596 |
| S1-Mid | DCIS | Development | 358,011 | 250,934 |
| S1-Bot | DCIS | Evaluation | 253,029 | 182,631 |
| S2-Top | Normal | Development | 279,734 | 228,291 |
| S2-Mid | DCIS | Evaluation | 348,804 | 282,581 |
| S2-Bot | DCIS | Evaluation | 336,019 | 257,392 |
| S3-Top | CCH/DCIS | Evaluation | 300,296 | 228,658 |
| S3-Mid | DCIS | Development | 371,809 | 277,809 |
| S3-Bot | DCIS | Evaluation | 148,878 | 105,203 |
| S4-Top | IDC | Evaluation | 187,107 | 134,274 |
| S4-Mid | IDC | Evaluation | 330,169 | 245,893 |
| S4-Bot | DCIS | Development | 200,942 | 139,927 |
| (B) ATERA breast tissue section |  |  |  |  |
| Characteristic | Description |  |  |  |
| Spatial transcriptomic platform | ATERA Whole Transcriptome Assay (10x Genomics) |  |  |  |
| Tissue preparation | FFPE |  |  |  |
| Diagnosis | Ductal carcinoma in situ (DCIS) |  |  |  |
| Receptor status | ER+/PR+/HER2– |  |  |  |
| Number of targeted genes | 18,028 |  |  |  |
| Allocation | External evaluation |  |  |  |
| Cell-type annotation source | 10x Genomics |  |  |  |
| Molecularly annotated cells | 168,326 |  |  |  |
| H&E nuclei with transferred labels | 115,904 |  |  |  |

The ATERA specimen was used exclusively for external evaluation and was not included in model training, validation, or hyperparameter selection. Cell-type annotations provided by 10x Genomics were harmonized into the seven cellular categories and transferred to H&E nuclear instances.
FFPE, formalin-fixed paraffin-embedded; DCIS, ductal carcinoma in situ; ER, estrogen receptor; PR, progesterone receptor; HER2, human epidermal growth factor receptor 2.
Xenium counts are from the final SingleR voting exports. H&E counts are from the whole-section label-transfer exports. Each H&E label was verified against the reconciled Xenium annotation for the corresponding cell ID.
Specimen diagnoses are from the 10x Genomics Xenium dataset.
Development sections contributed the following training and validation tile counts, respectively. S1-Mid, 59/10, S2-Top, 147/27, S3-Mid, 135/24, S4-Bot, 31/5.
CCH, columnar cell hyperplasia. DCIS, ductal carcinoma in situ. IDC, invasive ductal carcinoma.

**Table S2.** Single-cell references and cell-type harmonization. Related to Figure 2.

| Reference | Paper and public data records | Label field | Before QC | After QC |
|---|---|---|---|---|
| Wu *et al*. (2021) | Nature Genetics<br>GSE176078 · SCP1039 | celltype_minor | 100,064 | 98,418 |
| Dong *et al*. (2025) | Nature Communications<br>E-MTAB-14560 · E-MTAB-14566<br>CELLxGENE collection | Level2 | 10,689 | 8,853 |
| Chen *et al*. (2026) | NAR Genomics and Bioinformatics<br>CELLxGENE atlas | cell_type | 621,200 | 579,434 |
| Yan *et al*. (2026) | Nature<br>PRJNA1041570 · CELLxGENE collection | cell_type | 427,823 | 411,684 |
| **Reference** | **Harmonized category** | **Original labels in the harmonization code** |  |  |
| Wu *et al*. (2021) | Tumor | Cancer Basal SC • Cancer Cycling • Cancer Her2 SC • Cancer LumA SC • Cancer LumB SC |  |  |
| Wu *et al*. (2021) | Epithelial | Luminal Progenitors • Mature Luminal |  |  |
| Wu *et al*. (2021) | Myoepithelial | Myoepithelial |  |  |
| Wu *et al*. (2021) | Endothelial | Endothelial ACKR1 • Endothelial CXCL12 • Endothelial Lymphatic LYVE1 • Endothelial RGS5 |  |  |
| Wu *et al*. (2021) | Myeloid | Cycling_Myeloid • DCs • Macrophage • Monocyte |  |  |
| Wu *et al*. (2021) | Lymphocytes | B cells Memory • B cells Naive • Cycling T-cells • NK cells • NKT cells • Plasmablasts • T cells CD4+ • T cells CD8+ |  |  |
| Wu *et al*. (2021) | Stromal | CAFs MSC iCAF-like • CAFs myCAF-like • Cycling PVL • PVL Differentiated • PVL Immature |  |  |
| Dong *et al*. (2025) | Tumor | Tumor • Tu_B1 • Tu_B3 • Tu_B4 |  |  |
| Dong *et al*. (2025) | Endothelial | Vessel |  |  |
| Dong *et al*. (2025) | Myeloid | Mast_cell • Myeloid |  |  |
| Dong *et al*. (2025) | Lymphocytes | B • T_NK |  |  |
| Dong *et al*. (2025) | Stromal | Fibro_muscle |  |  |
| Chen *et al*. (2026) | Tumor | malignant cell |  |  |
| Chen *et al*. (2026) | Endothelial | capillary endothelial cell • endothelial cell • endothelial cell of artery • endothelial cell of vascular tree • vein endothelial cell |  |  |
| Chen *et al*. (2026) | Myeloid | conventional dendritic cell • cycling macrophage • macrophage • mast cell • myeloid dendritic cell • plasmacytoid dendritic cell |  |  |
| Chen *et al*. (2026) | Lymphocytes | CD4-positive helper T cell • CD4-positive, CD25-positive, CCR4-positive, alpha-beta regulatory T cell • CD8-positive, alpha-beta regulatory T cell • IgG plasma cell • T follicular helper cell • cycling T cell • effector memory CD4-positive, alpha-beta T cell • effector memory CD8-positive, alpha-beta T cell • exhausted T cell • memory B cell • naive T cell • natural killer cell |  |  |
| Chen *et al*. (2026) | Stromal | cycling stromal cell • fibroblast • myofibroblast cell • pericyte • vascular associated smooth muscle cell |  |  |
| Yan *et al*. (2026) | Tumor | abnormal cell |  |  |
| Yan *et al*. (2026) | Endothelial | endothelial cell |  |  |
| Yan *et al*. (2026) | Myeloid | myeloid leukocyte |  |  |
| Yan *et al*. (2026) | Lymphocytes | T cell • lymphocyte of B lineage |  |  |
| Yan *et al*. (2026) | Stromal | fibroblast • perivascular cell |  |  |

Counts are from the reference quality-control reports and precede per-label capping.
Original reference labels are reproduced verbatim. Only the Wu reference supplied labels that mapped to the epithelial and myoepithelial categories.

**Table S3.**
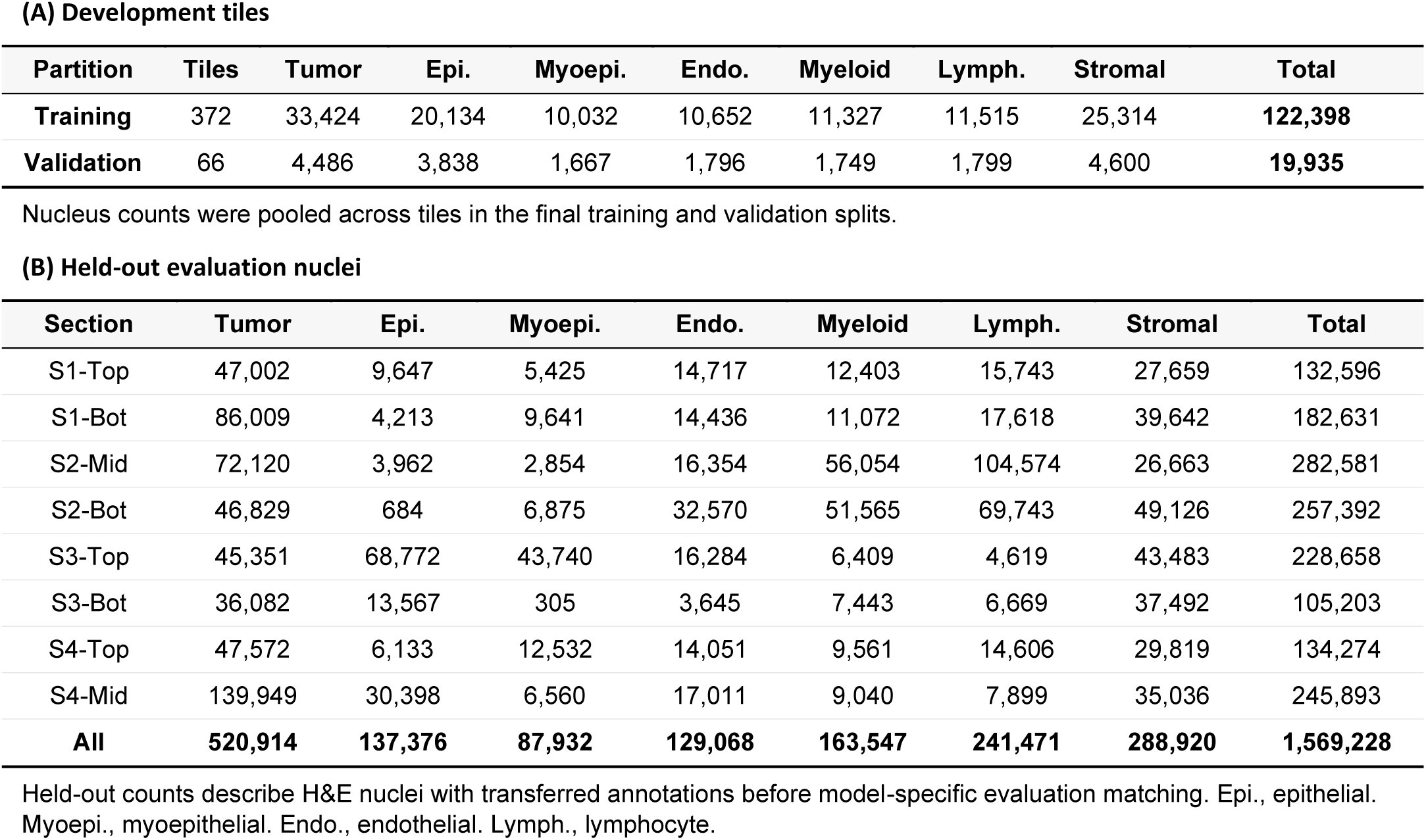
Cell-class distributions in development and evaluation data. Related to Figures 3 and 4.

| (A) Development tiles |  |  |  |  |  |  |  |  |  |
|---|---|---|---|---|---|---|---|---|---|
| **Partition** | **Tiles** | **Tumor** | **Epi.** | **Myoepi.** | **Endo.** | **Myeloid** | **Lymph.** | **Stromal** | **Total** |
| **Training** | 372 | 33,424 | 20,134 | 10,032 | 10,652 | 11,327 | 11,515 | 25,314 | **122,398** |
| **Validation** | 66 | 4,486 | 3,838 | 1,667 | 1,796 | 1,749 | 1,799 | 4,600 | **19,935** |
| **(B) Held-out evaluation nuclei** |  |  |  |  |  |  |  |  |  |
| **Section** | **Tumor** | **Epi.** | **Myoepi.** | **Endo.** | **Myeloid** | **Lymph.** | **Stromal** | **Total** |  |
| S1-Top | 47,002 | 9,647 | 5,425 | 14,717 | 12,403 | 15,743 | 27,659 | 132,596 |  |
| S1-Bot | 86,009 | 4,213 | 9,641 | 14,436 | 11,072 | 17,618 | 39,642 | 182,631 |  |
| S2-Mid | 72,120 | 3,962 | 2,854 | 16,354 | 56,054 | 104,574 | 26,663 | 282,581 |  |
| S2-Bot | 46,829 | 684 | 6,875 | 32,570 | 51,565 | 69,743 | 49,126 | 257,392 |  |
| S3-Top | 45,351 | 68,772 | 43,740 | 16,284 | 6,409 | 4,619 | 43,483 | 228,658 |  |
| S3-Bot | 36,082 | 13,567 | 305 | 3,645 | 7,443 | 6,669 | 37,492 | 105,203 |  |
| S4-Top | 47,572 | 6,133 | 12,532 | 14,051 | 9,561 | 14,606 | 29,819 | 134,274 |  |
| S4-Mid | 139,949 | 30,398 | 6,560 | 17,011 | 9,040 | 7,899 | 35,036 | 245,893 |  |
| **All** | **520,914** | **137,376** | **87,932** | **129,068** | **163,547** | **241,471** | **288,920** | **1,569,228** |  |

Nucleus counts were pooled across tiles in the final training and validation splits.
Held-out counts describe H&E nuclei with transferred annotations before model-specific evaluation matching. Epi., epithelial. Myoepi., myoepithelial. Endo., endothelial. Lymph., lymphocyte.

## Acknowledgements

This work was supported by the following sources: U54CA268083, UG3CA275681, all from the NIH; the Break Through Cancer Data Science TeamLab; and the Johns Hopkins University and Instituto Superior Técnico student exchange program (HOPTEC and FLAD). D.F.V. acknowledges financial support from the FCT PhD Studentship 2024.00637.BD.

## Author contributions

All authors conceived and designed the study. D.F.F., D.F.V., V.Q. and M.W. developed the methodology and implemented the computational framework. D.F.F., D.F.V., M.W. and V.Q. performed the experiments, processed the data and conducted the formal analyses. D.F.F., D.F.V., M.W., V.Q. and S.J. contributed to data acquisition, annotation and curation. D.F.F., D.F.V., V.Q., M.W., A.F. and A.K. interpreted the results. D.F.F., D.F.V., and V.Q. prepared the figures and wrote the original draft of the manuscript. A.K., A.F., P.W. and D.W. supervised the study, provided resources and acquired funding. All authors contributed to reviewing and editing the manuscript and approved the final version.

## Additional information

The authors declare no competing interests.

